# A Moving Target: The Megaplasmid pMPPla107 Sensitizes Cells to an Inhibitory Agent Conserved Across *Pseudomonas spp*

**DOI:** 10.1101/537589

**Authors:** Brian A. Smith, Yelena Feinstein, Meara Clark, David A. Baltrus

## Abstract

The widespread use of antibiotics across clinical and agricultural settings results in strong selection pressures and contributes to the fixation of antibiotic resistance genes, the presence of which lowers the efficacy of proven treatments for infection. Furthermore, plasmids are often key vectors that facilitate the rapid dispersal of antibiotic resistance genes across bacterial strains via horizontal gene transfer. In contrast to previous widespread correlations between plasmid acquisition and resistance to antimicrobial compounds, we demonstrate that acquisition of the *P. syringae* megaplasmid pMPPla107 sensitizes strains to a conserved, bacteriostatic small molecule produced by many *Pseudomonas spp*. Moreover, we find the acquisition of pMPPla107 reduces production of the inhibitory agent. Our results provide insights and suggest new directions to investigate collateral sensitivity to antimicrobials due to plasmid acquisition as well as highlight costs associated to horizontal gene transfer in the form of sensitivity to antagonistic microbial interactions.

## INTRODUCTION

Bacteria exist in complex ecological networks where interactions between community members range from mutualistic to antagonistic. Individual strains have therefore developed a variety of methods to interact with and/or outcompete surrounding bacterial cells including the production of secondary metabolites that can inhibit or allow growth of various microorganisms, as well as bacteriocins that kill target cells with high specificity(1–9). Secondary metabolites can also provide neighboring microbes a previously missing metabolic resource allowing for colonization of previously uninhabitable niches (10, 11). Such interactions play a critical role in the development and maintenance of bacterial communities under natural conditions, but the discovery and exploitation of antimicrobials underlying these interactions has transformed the management of infection under both clinical and agricultural settings.

Horizontal gene transfer (HGT) allows for the alteration of genotypes by the rapid acquisition of genes via mobile genetic elements like plasmids and phage. Although acquisition of plasmids is commonly associated with beneficial effects in recipient strains, like the colonization of new niches through genes associated with symbiosis, virulence, and antibiotic resistance, numerous costs of plasmid acquisition have also been identified. Phenotypic costs are typically assumed to be driven by draining of limiting resources within the cell by plasmid borne pathways (e.g. use of ATP and nucleotides to promote plasmid replication) or through costly protein interactions that disrupt cellular networks(12–14), which both manifest as lower overall fitness under permissive conditions in cells containing plasmids compared to those that are plasmid free. Along with intracellular costs of plasmids, creation of the type IV secretion pilus enables killing of conjugating donor cells by phage and type VI secretion systems(12, 15). These various costs highlight the fact that most plasmids can be considered as selfish parasites of their host cells, regardless of the potential benefits they may provide, and that the existence of these costs may prove to be an Achilles heel that can be exploited for the development of novel treatments to limit the spread of antibiotic resistance genes (or any other type of gene) that can be disseminated through plasmid conjugation.

*Pseudomonas* is a diverse genus of Gram-negative bacteria that includes a variety of potential pathogens like *P. syringae* and *P. aeruginosa. P. syringae* is a phytopathogen that causes economic loss in a wide variety of fruits, vegetables, and ornamental plants across the globe(16–18). The second species, *P. aeruginosa* is an opportunistic pathogen of humans with multiple drug resistances that is a main driver in morbidity and mortality in cystic fibrosis patients, burn victims, and the immunocompromised(19, 20). Therefore, identifying novel methods to limit infection by these pathogens could have dramatic impacts on clinical and agricultural settings. Previously we found that the *Pseudomonas syringae* megaplasmid pMPPla107 leads to numerous phenotypic costs when acquired by alternative hosts including: thermosensitivity, alterations in biofilm production and motility, and collateral sensitivity to some antibiotics (21). Understanding how HGT can create these costly effects may enable new methods for clinical and agricultural treatments of pathogens as well as a better understanding of HGT dynamics among plasmids in general.

In addition to these previously identified phenotypic costs, we have demonstrated that acquisition of pMPPla107 sensitizes strains to growth inhibition by an unknown compound found within the supernatant of P. aeruginosa PA14. Here we follow up on our previous publication and characterize the inhibitory phenotype associated with pMPPla107 and the supernatant of *P. aeruginosa* (21). We optimize assays for both the timing of production of and sensitivity to this inhibitory agent, identifying organisms that produce and/or are susceptible to this inhibitory agent, and screen to demonstrate that no mutant present within the *P. aeruginosa* PA14 transposon mutant library tested eliminates production of this compound. Fluorescence microscopy experiments further reveal that cells were not lysed when treated with the inhibitory agent, indicating the compound has bacteriostatic properties. Using liquid chromatography-mass spectroscopy (LC-MS) we identify a mass of interest with a fraction with growth inhibition properties. This body of work demonstrates a unique system where a large mobile element sensitizes cells to a common molecule produced by Pseudomonads and provides further evidence of widespread phenotypic costs that accompany HGT.

## METHODS

### Collection of Supernatants

All strains used in this report are listed in Table 1 (DOI: <u>doi.org/10.6084/m9.figshare.7627334.v1</u>). Pseudomonas strains and *E. coli* were grown in various media types and used in this work for the collection of supernatants (Table 1). Additionally, these Pseudomonas strains (Table 1) were grown on bacterial overlays. Although these strains carry various antibiotic resistance loci, antibiotics were not used in the bacterial overlay assays to reduce unwanted inhibitory affects from antibiotics.

**Table 1:** Strain list containing information for all the strains used in this manuscript. File is available at Figshare (DOI: <u>doi.org/10.6084/m9.figshare.7627334.v1</u>)

To collect supernatants single colonies were picked from agar plates containing the strain’s preferred media type to 2mL of the organism’s preferred media type overnight with shaking (200rpm) at either 27°C or 37°C dependent on the organism (Table 1). Media types used were Salt Water-LB (SW-LB), Lysogeny broth (LB), or King’s medium B (KB) and their recipes can be found at: dx.doi.org/10.17504/protocols.io.xf3fjqn. Cells were then diluted 1:100 to fresh media in the same media type previously used and allowed to grow for either 24 or 48 hours (Table 1) with shaking (200rpm) at preferred 27°C or 37°C (unless specified) to reach saturation based off previous growth curves(22) and turbidity. Saturated cells were then centrifuged at 10,000 × *g* for 5 minutes. Supernatants were collected and were filter sterilized by passing samples through a 0.22μm filter using a syringe. Samples were stored at 4°C.

### Bacteriocin Isolation

Bacteriocin induction and isolation was carried out as per Hockett *et al* (7). *P. syringae* B15 cultures were made in 3mL of KB with shaking (200rpm) at 27°C for 24 h and were then diluted 1:100 to fresh KB media the next day. Cultures were allowed to grow for approximately 4 h or until slight turbidity was visualized and at which point mitomycin C was added to a final concentration of 0.5μg/mL. Cultures were then allowed to grow over night and were filter sterilized with a 0.22μm filter and stored at 4°C.

### Physical Characterization of the Inhibitory Agent

Filter sterilized supernatants containing the inhibitory agent or isolated bacteriocins were treated with proteinase K (proK), autoclave, or precipitated with PEG 8000 to determine the presence of proteins. For proK treatment, the inhibitory agent or isolated bacteriocins were mixed with proK to a final concentration of 1.0mg/mL (Invitrogen) and mixtures were incubated at 37°C for 2 h. ProK was then heat inactivated by incubation of samples at 95°C for 10 min. The inhibitory agent and bacteriocins were heat treated by autoclaving samples at 121°C for 15 min at 30 psia. Concentration of larger molecules within supernatants followed a polyethylene glycol (PEG) precipitation protocol previously published by Hockett *et al*(7). Briefly, samples are mixed with NaCl (to a 1M final concentration), incubated on ice for 30min, mixed with PEG 8000 (10% final concentration) and left on ice for 1 h, centrifuged at 16,000 × *g* for 30 min at 4°C, and pellets are suspended in buffer (0.01 Tris, 0.01 M MgSO4, pH 7) at 0.1 or 0.01 the original volume.

### Bacterial Overlay

A single colony was picked from an agar plate containing the strain’s preferred media type and allowed to grow overnight in 2mL of preferred media (SW-LB, KB, LB) at 27°C with shaking. The following day cells were diluted 1:100 (30μL cells:2.97mL media) into fresh media and allowed to grow for 48 h at 27°C with shaking and 200rpm. After 48 h of cell growth, all cultures are diluted 1:100 (30μL cells:2.97mL media) to fresh KB and allowed to grow for 4 h at 27°C with shaking. Cultures are then mixed 1:10 (300μL cells:2.7mL agar) with molten 0.4% agar and poured onto a KB agar plate. Plates were allowed to solidify for 15 min, at which time 10 μL spots of either the inhibitory agent or the bacteriocin were spotted onto the agar. Spots were allowed to dry and plates were incubated at 27°C for 16-24 h.

### Bacterial Conjugations With pMPPla107

Biparental bacterial conjugations were performed using a *P. syringae* (DBL328) pMPPla107 and a recipient strain, either DBL73 or DBL306, in a 1:1 ratio (750μL:750μL). Cells were mixed and centrifuged at 3,000 × *g* for 3min and supernatants were removed by decanting without disturbing the pellet. Pellets were washed and reconstituted in 1mL of 10mM MgCl_2_. Centrifugation and washing steps were repeated. 10μL and 100μL samples of reconstituted sample were spread to a KB plates containing selection for pMPPla107 and the recipient strain using either Rif50μg/mL and Kan30μg/mL (DBL73) or Tet10μg/mL, Cm30μg/mL, and Kan30μg/mL (DBL306) and grown for 24-48hr at 27°C. Resistant colonies were grown in liquid culture containing the same antibiotic combination and using diagnostic primers(21) PCR was performed for the presence of pMPPla107. Resistant cells that resulted in amplification were preserved at −80°C in 40% glycerol.

### Liquid Chromatography

After 48hr of growth, *Pseudomonas stutzeri* 28a24 (DBL408)(21) supernatants prepared as above were separated via Agilent Zorbax 300 SB-C18, 5um, 250 x 2.10mm column using Paradigm MS4B - multi-dimensional HPLC separations module (Michrom BioResources, Inc., Auburn, Calif.) with UV detection @220nm via WellChrom spectrophotometer K-2501 (Knauer; Berlin, Germany) inline with Gilson FC 203B fraction collector (Gilson, Inc., Middleton, WI). The mobile phases 0.1%Formic acid (FA)/H2O (A) and 0.1% FA/ACN (B) were delivered at a flow rate of 200 μl/min with the following gradient: 5%B (0-2.0 min), from 5 to 100% B (2.0-30.0 min), 100%B (30.0-35.0 min), from 100 to 5%B (35.0-35.2 min), 5%B (35.2-40 min).

6ul of sample/per injection were fractionated into 1.5min fractions, the fractions were dried down in a vacuum centrifuge, reconstituted in 10μL of sterile H_2_O and spotted onto overlay plates to test for the presence of inhibitory activity against pMPPla107 (DBL408) as per above.

### Low-resolution Mass Spectrometric (MS) Analysis

MS analysis of supernatant fractions collected from liquid chromatography was performed using Bruker Daltonics amaZon SL Ion Trap mass spectrometer (Bruker Daltonics Inc, Billerica, MA) by direct infusion. The fractions were vacuum centrifuged until dry and reconstituted in 10μL of H_2_O and tested for inhibitory activity against pMPPla107 (DBL408) on an overlay plate as previously described. LC-MS analysis of the analyte was performed via amaZon as well with the sample introduction via the ThermoFisher Scientific UltiMate 3000 HPLC system using the same column and the gradient as described in “Liquid chromatography” section.

### Replication and screening of the P. aeruginosa PA14 Transposon Mutant Library

*P. aeruginosa* PA14 transposon mutant library plates from the Ausubel Lab at Harvard(23) were replicated using LB with the suggested antibiotic according to the manual (http://pa14.mgh.harvard.edu/pa14/downloads/manual.pdf) was added to three round-bottomed 96 well micro titer plates using a peristaltic pump. Plates were inoculated and grown at 37°C and 100% humidity with shaking to eliminate evaporative loss. Humidity was measured using a hygrometer. After 24 hours of growth 50μL of 80% glycerol was added to all daughter replicate plates. Replicate plates were sealed with sterile aluminum covers and transferred to a Beckman Coulter Biomek FX to mix glycerol by shaking of plates for 1 min. Replicate plates were frozen at −80°C.

Library plates were revived in 96 well micro titer plates with 175μL of LB with gentamicin (15ng/μL) for the *MAR2xT7* transposon, or kanamycin (30ng/μL) for the *PhoA* transposon for 24 hours at 37°C with shaking at 100% humidity. 75μL were transferred to a 96 well polypropylene PCR plate (TempPlate, USA Scientific) and sealed with a sterile aluminum seal (Aluma Seal II, Excel Scientific). Sealed plates were sterilized by heating cells to 95°C for 10min in a thermocycler. 5μL from each well was spotted onto an overlay plate with cells containing pMPPla107 (DBL494). Overlay plates followed a similar protocol as above, except that a larger petri dish (150mm x 50mm) was used and requires 900μL of cells mixed with 8.1mL of molten agar.

### Fluorescence Microscopy

Following the overlay protocol above, cells grown for 4hr in KB were than mixed 1:1 with *P. stutzeri* supernatant or a negative control (KB) for 15min. Afterward 3μL of the mixed LIVE/DEAD kit (L7012, Invitrogen) was added to the samples and mixed thoroughly. Samples were placed on a CELLview TC treated, one compartment, sterile culture dish (Greiner) and viewed under a Lecia DMI6000 inverted microscope. Images were captured using a Hamamatsu Flash 4.0 and Lecia LAS-X software.

## Results

### Sensitivity is Dependent on Media Type and Growth State

To ensure reprocibility across numerous assays, we first sought to characterize and optimize sensitivity to the previously identified inhibitory agent. Using strain DBL408 as an indicator and strain DBL736 as a producer, we found that both media type and time to plating are critical for successful overlays. Inhibitory activities of *P. aeruginosa* supernatants appear more distinct and clear against strain DB408 when plating overlays onto KB compared to LB agar (Figure 1) and we therefore decided to use KB media for overlays of the target strains.

**Figure 1:**
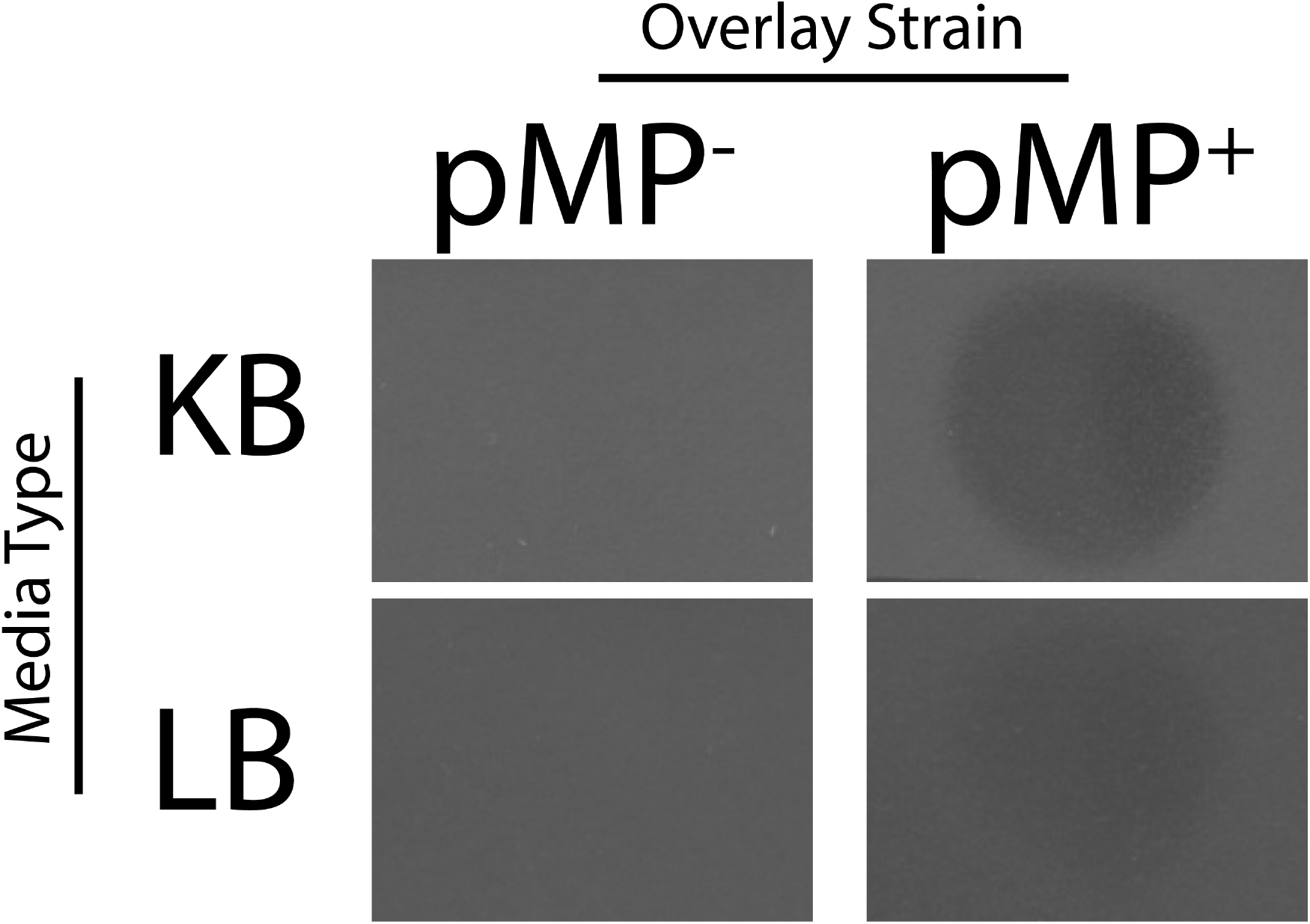
Media Type Affects Clarity of Inhibitory Phenotype on Bacterial Overlays. *P. stutzeri* with (DBL408) and without (DBL386) the megaplasmid were grown in either LB and KB for four hours, mixed with molten agar, and poured onto a KB or LB plate. 10μL spots of filter sterilized stationary phase supernatants of *P. aeruginosa* PA14 were then spotted on to overlays. Zones of inhibition can be seen for both overlays containing pMPPla107 although KB media causes the phenotype to be more distinct. pMP^−^ = no megaplasmid, pMP^+^ = pMPPla107. Images are representative of at least 3 replicates.

Because antibacterial sensitivity is increased during log phase we also tested different growth times when plating target strains to an overlay to optimize efficacy of the inhibitory agent(24, 25). After cells were grown with shaking to saturation, they were diluted 1:100 (cells:media) and plated after 2, 3, 4, and 5 hours of growth at 27°C. Our results indicate that strains that lack pMPPla107 show no difference in sensitivity to the supernatant between all time points in the absence of pMPPla107, while strains containing the megaplasmid have distinct zones of inhibition at 4 and 5 hours of growth while hours 2 and 3 have weaker activity (Figure 2). However, we further note that inhibitory activity could no longer be clearly visualized if cells were grown for longer than 5 hours (Data not shown). Therefore, for the remainder of assays reported here, strains were grown for 4 hours prior to overlay plating unless specified.

**Figure 2:**
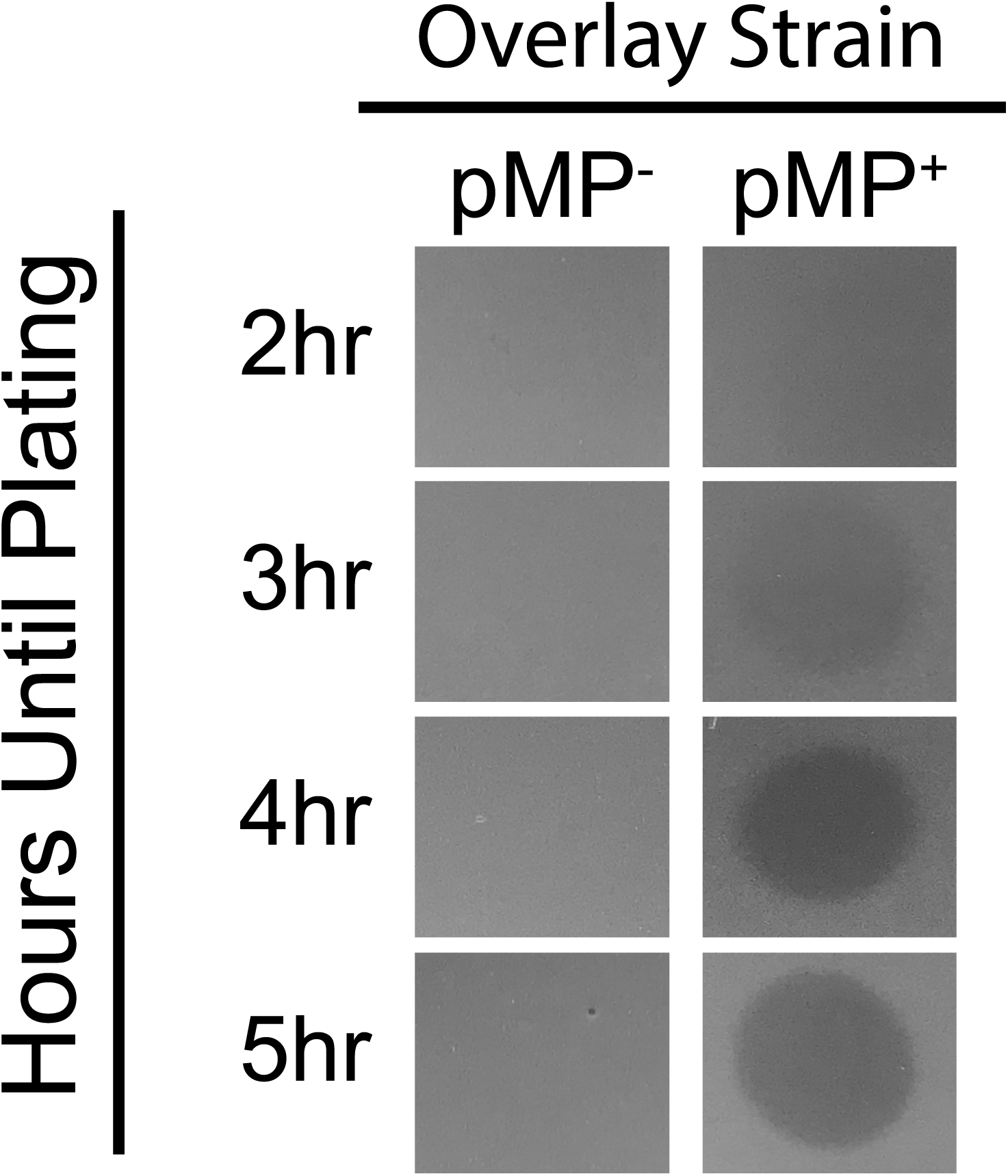
The optimum time to plate strains to an overlay occurs at hour 4. Overlay strains were allowed to grow for 2, 3, 4, and 5hr after a 1:100 dilution was made into KB media. 10μL spots of filter sterilized stationary phase supernatants of *P. aeruginosa* PA14 Distinct clearing zones are present at 4 hours. pMP^−^ = no megaplasmid, pMP^+^ = pMPPla107. Images are representative of at least 3 replicates.

### Production of the Inhibitory Agent Correlates With P. stutzeri Growth

To optimize collection of the inhibitory agent, we collected supernatants over a range of time points during growth of the producer strain. Production of the inhibitory agent occurs within log phase around hour 6(26). We found that inhibitory activity increases at hours 12 and 16, but that inhibition from 16 hour old cultures appears similar to that at 24 and 48 hours (Figure 3). Given these results we collect filter-sterilized supernatants between hours 24-48 from this point on unless otherwise specified.

**Figure 3:**
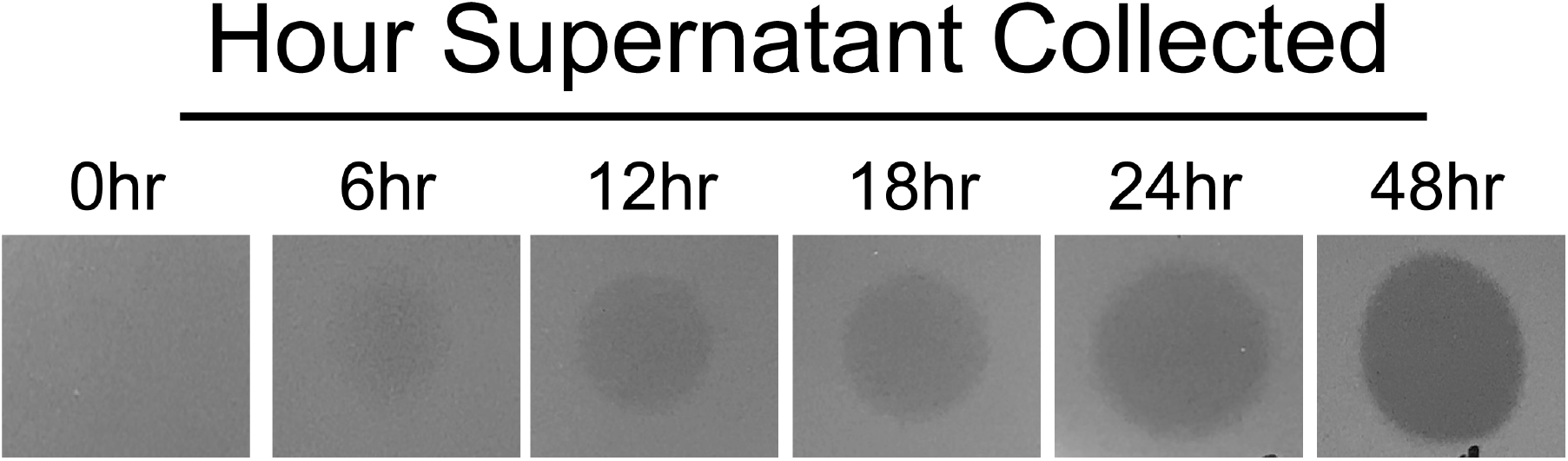
Inhibitory Activity in pseudomonas supernatants increases with growth after 6 hours. Filter sterilized supernatants were collected from *P. stutzeri* (DBL386) at 0, 6, 12, 18, 24, and 48 hours after a 1:100 dilution into SW-LB. 10μL of these various time points were spotted onto an overlay of *P. stutzeri* with pMPPla107 (DBL408) Images are representative of at least 3 replicates.

### Acquisition of pMPPla107 Sensitizes Strains Most Across the Pseudomonas Phylogeny to the Inhibitory Agent

Previously the only strain tested for sensitivity to the *P. aeruginosa* supernatants was DBL408 (DBL386+pMPPla107) and DBL386, yet there is also data suggesting that pMPPla107 can conjugate to other *Pseudomonas* spp.(21, 26). Therefore, we tested whether sensitization to the inhibitory agent by acquisition of pMPPla017 varied across the *Pseudomonas* phylogeny (Figure 4). According to these assays, we found that sensitization is present if pMPPla107 is acquired by *P. stutzeri, P. syringae*, and *P. fluorescens*. However, the sensitization phenotype appears to be absent in *P.putida*.

**Figure 4:**
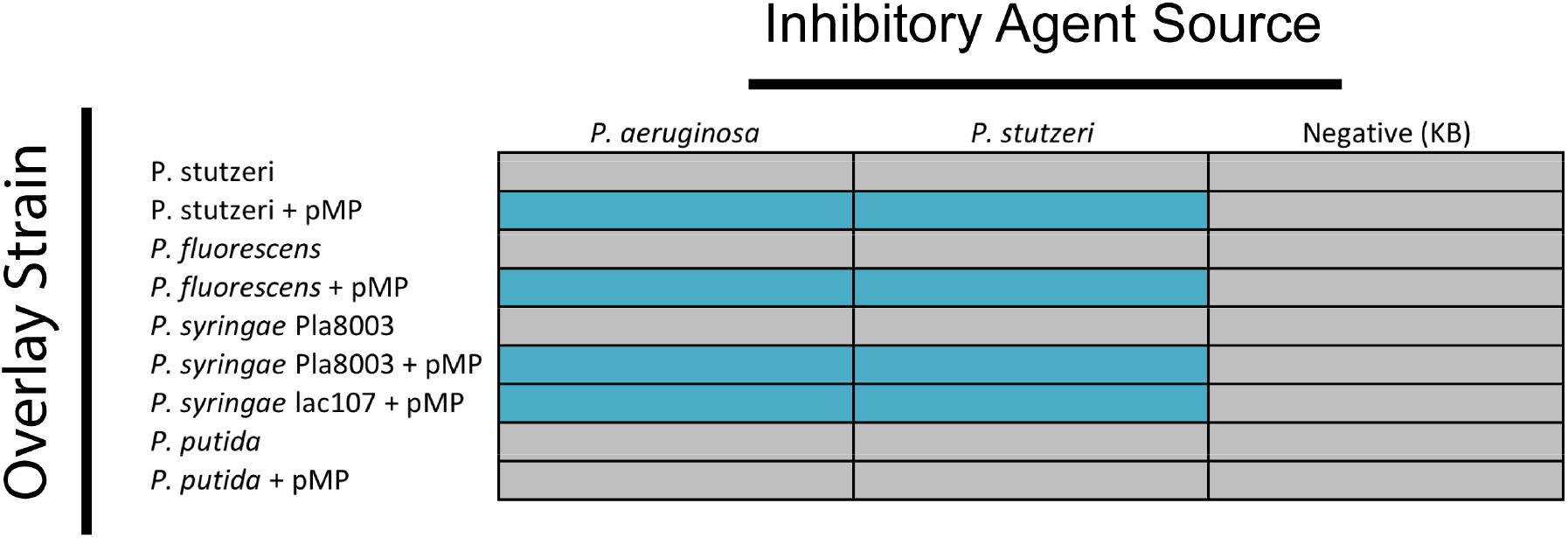
pMPPla107 acquisition causes sensitivity to the inhibitory agent in various *Pseudomonas spp*. We tested the inhibitory agent on various overlays of *Pseudomonas spp*. with and without pMPPla107. Blue = inhibited, grey = no inhibition, pMP = pMPPla107. Data are representative of at least 3 replicates.

### The Inhibitory Agent is Not a Protein

Bacteria utilize various methods to inhibit the growth of neighboring microorganisms that include both proteins and small molecules. To determine the molecular nature of the inhibitory agent we pulled down *P. aeruginosa* supernatants with PEG 8000, treated supernatants with proK, and autoclaved supernatants. As a protein control for treatments with proK and heat we isolated bacteriocins from *P. syringae* pv. B15 which has known killing activity of *P. syringae* pv. *morsprunorum*(7). Treatment with proK and autoclaving eliminates activity of the *P. syringae* bacteriocin, but the inhibitory agent that targets pMPPla107 remains active (Figure 5). Furthermore, PEG 8000 is used to precipitate and concentrate large molecules including proteins, but our results indicated that activity was reduced after PEG precipitation (Supplemental Figure 1). Together these data indicate that the inhibitory agent, which specifically targets pMPPla107, is not a protein and is likely to be a relatively small molecule.

**Figure 5:**
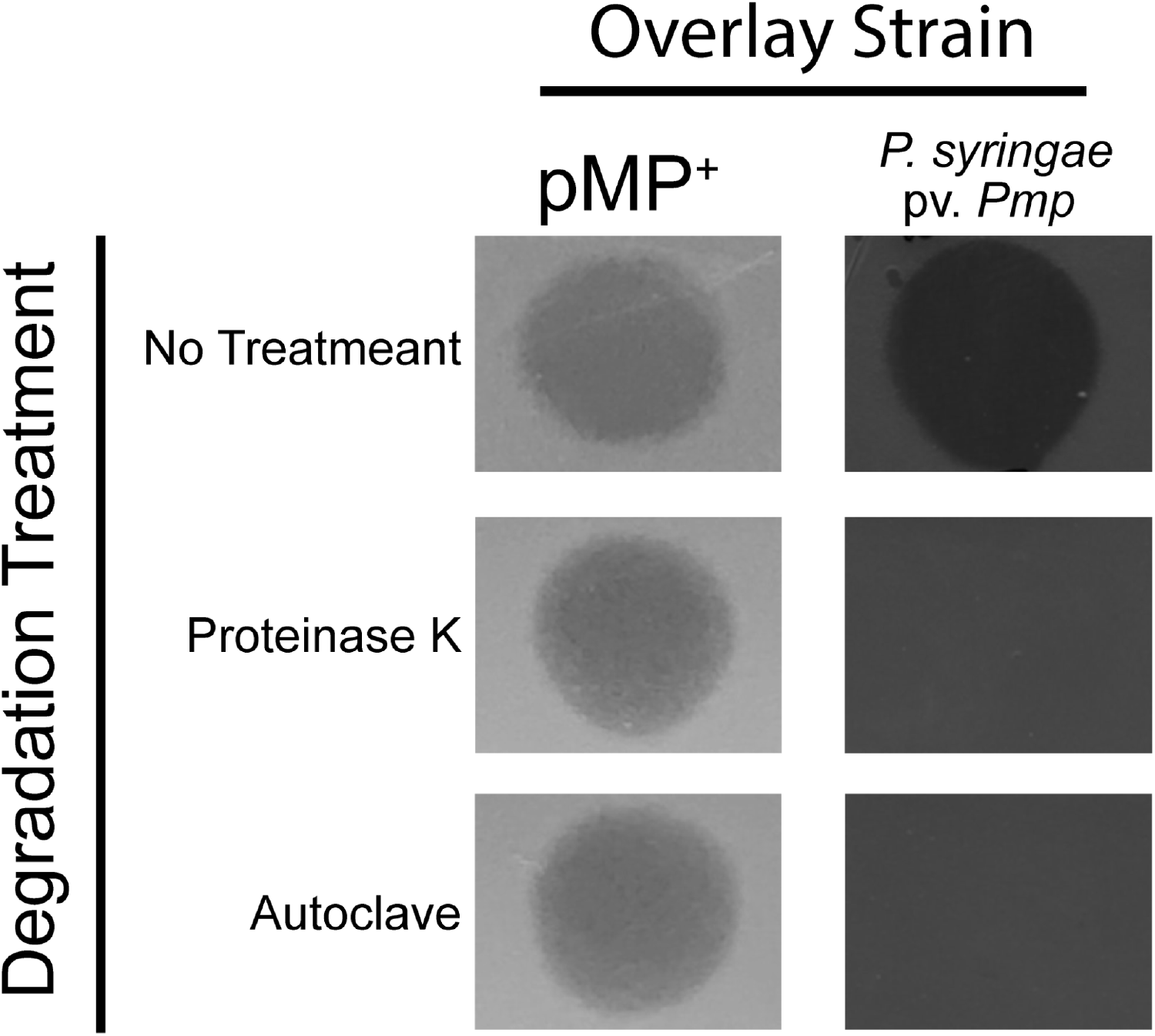
Protein degradation treatments do not eliminate activity of the inhibitory agent. *P. stutzeri* with pMPPla107 (DBL408) were spotted with 10μL of filter sterilized supernatants from *P. aeruginosa* PA14, and *P. syringae* pv. *morsprunorum* (*Pmp*) were spotted with 10μL of bacteriocins isolated from *P. syringae* B15. Treatments against the inhibitory agent and B15 bacteriocin consisted of either Proteinase K 1.0mg/mL incubated at 37°C for 2hrs or heat treated by autoclave at 121°C for 15min. pMP^+^ = *P. stutzeri* with pMPPla107. Images are representative of at least 3 replicates.

### Identification of a 530.70 m/z With Inhibitory Activity Against pMPPla107

We fractionated raw supernatants to identify the molecule responsible for the inhibitory activity using liquid chromatography and mass spectroscopy. Although UV chromatograms revealed that the samples were complex, 3 fractions demonstrated inhibition of pMPPla107 (Figures 6A-B). MS analysis of fraction 1 revealed a distinct peak occurring at a mass of 530.72 in positive mode and 528.70 in negative mode indicating a potential mass of 530.70 for the inhibitory agent. We were further able to determine a retention time of 15.5min using LCMS (Figure 6C Supplemental Figures 2 and 3). This retention time allowed us to eliminate various peaks from fraction 1 as LCMS analysis revealed peaks only present for masses of 513, 530, and 551 with retention times of 15.5min (Supplemental Figure 4). However, due to low concentrations of the suspected molecule we were unable to isolate enough sample for structural analysis with NMR.

**Figure 6:**
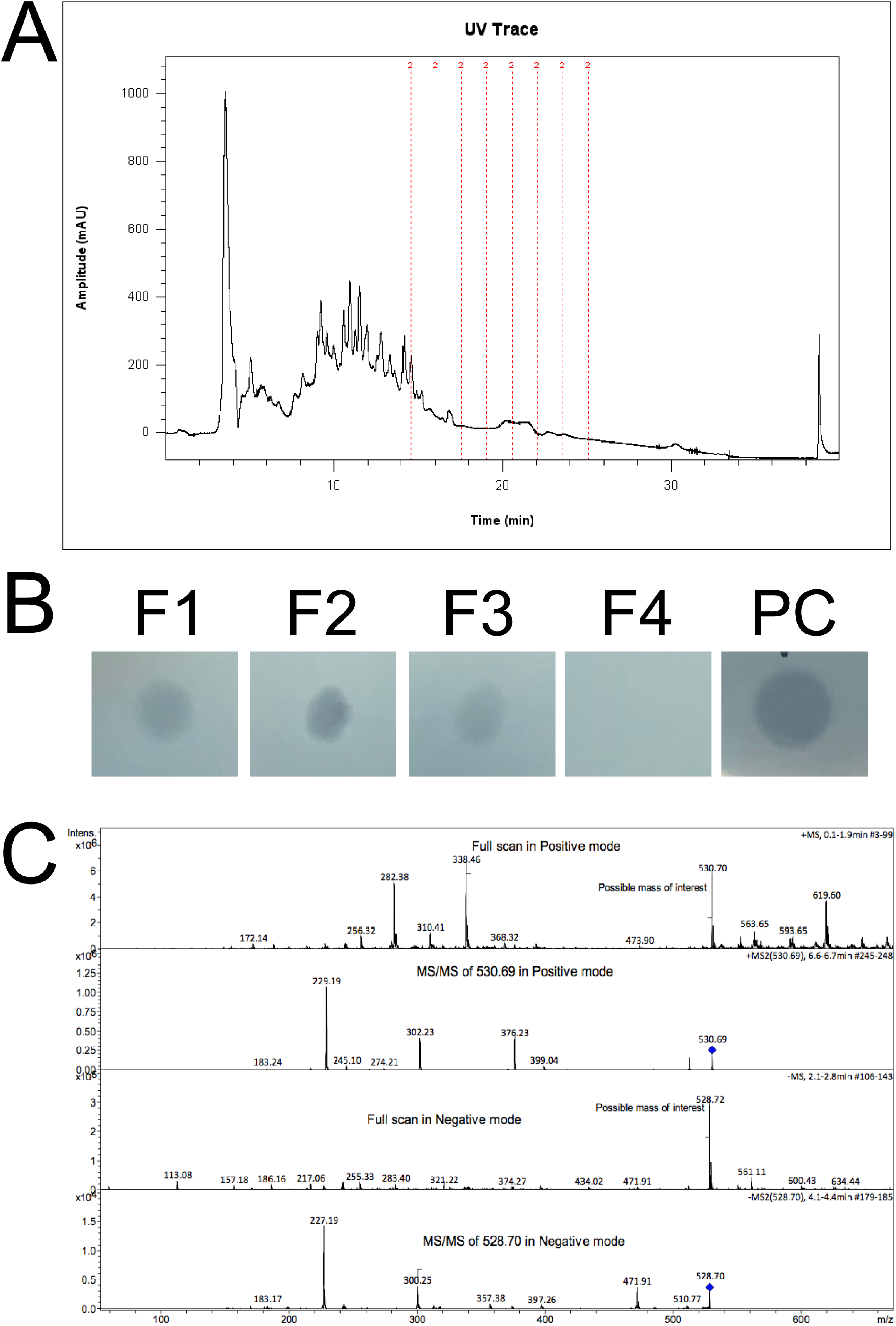
The MS spectrum of the active fraction via low resolution Mass Spectrometric Analysis. **A)** UV Chromatogram of 6μL *P. stutzeri* filter sterilized supernatants fractioned with 1.5min intervals collected between 14.5 and 25.5min (Red lines) indicates. **B)** 10μL spots of active fractions 1, 2, 3, (F1, F2, F3) and *P. stutzeri* supernatant (PC = positive control) on an overlay plate containing pMPPla107. **C)** MS analysis of fraction 1 was performed using Bruker Daltonics amaZon SL Ion Trap mass spectrometer (Bruker Daltonics Inc, Billerica, MA) by direct infusion. There is an abundant 530.72 mass observed in Positive mode and 528.70 - in Negative mode indicating a potential mass of 530.70. MS/MS indicates fragmentation of the peak with a blue diamond.

### No Mutants Tested from the PA14 Mutant Library Eliminate Production of the Inhibitory Agent

To identify a potential genetic basis for production of the inhibitory agent we replicated and developed a high throughput screening method of the entire *P. aeruginosa* PA14 non-redundant transposon mutant library(23). After isolating supernatants and screening 5,364/5,459 mutants against *P. stutzeri* with pMPPla107 (DBL494), we found that none of the transposon mutants eliminated or reduced activity against DBL494 (Table 2 DOI: <u>doi.org/10.6084/m9.figshare.7390235.v1</u> and Supplemental Figure DOI: 10.6084/m9.figshare.7631585). 95 mutants were missing from the library, as these were lost in the transfer and replication process.

**Table 2:** No knockouts in the tested PA14 knockout library stop or reduce production of the inhibitory agent. File is available at Figshare (DOI: <u>doi.org/10.6084/m9.figshare.7390235.v1</u>)

### Multiple Pseudomonas Species’ Supernatants Display Inhibitory Activity

Since the transposon screen suggests that production of the inhibitory agent is potentially an essential gene in *P. aeruginosa*, we hypothesized that this gene or pathway may be conserved across Pseudomonads. Therefore, we collected supernatants from various isolates across the pseudomonas phylogeny and found that production of the inhibitory agent is indeed conserved when tested against DBL408 (Figure 7). However, we also observed variations in production capabilities of different species. For instance, *P. putida* and *P. fluorescens* appear to produce the inhibitory agent at higher concentrations or produce a secondary killing agent that inhibits growth of non-megaplasmid strains (Figure 7). *P. fluorescens* supernatants are still more effective against strains containing pMPPla107 than those lacking pMPPla107 indicating that the megaplasmid still demonstrates sensitizing effects to the stationary phase supernatants of Pseudomonads (Figure 7).

**Figure 7:**
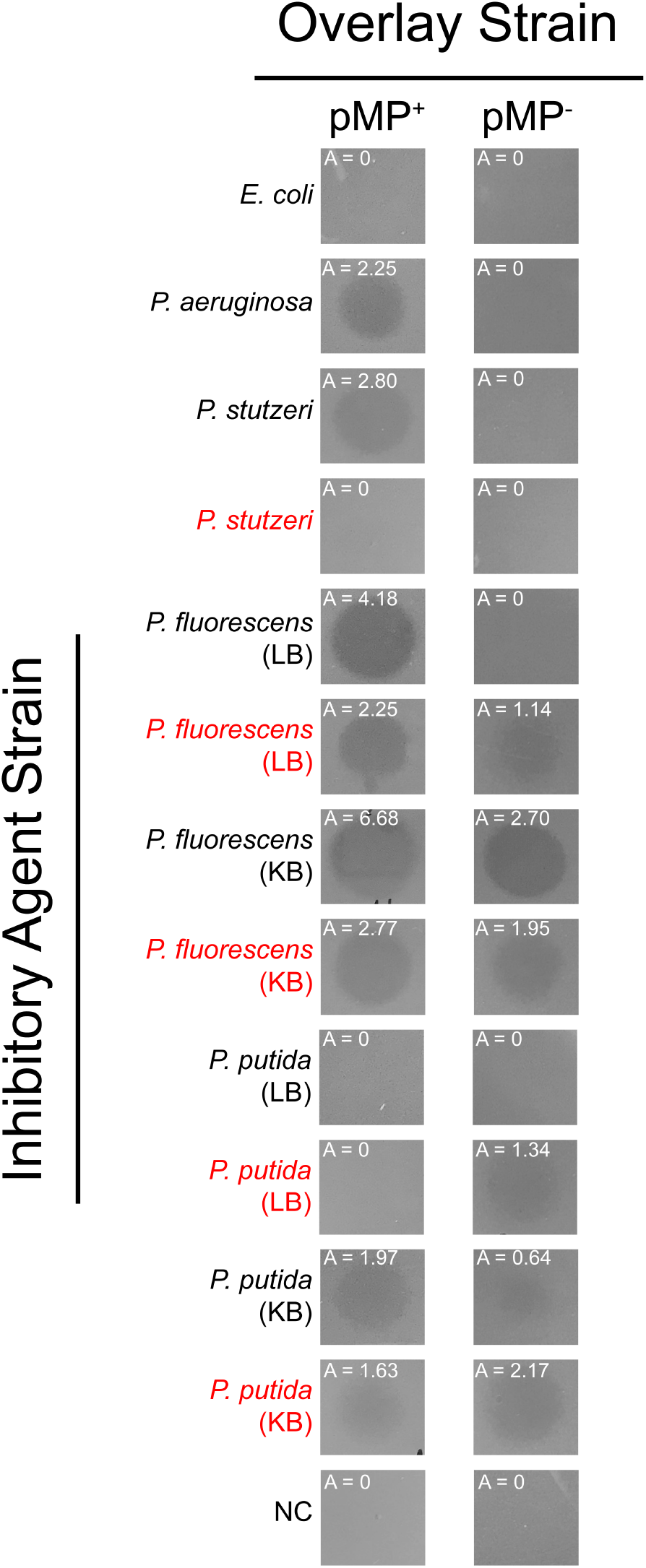
The inhibitory agent is conserved across the pseudomonas phylogeny and acquisition of pMPPla107 reduces production of the inhibitory agent. Supernatants were collected from various *Pseudomonas spp*. at stationary phase and tested for production of the inhibitory agent. Cells were grown in KB and LB before supernatants were collected to identify potential differences in production due to differences in metabolism. Black lettering = no megaplasmid, red lettering = pMPPla107, NC = negative control (10μL of KB). pMP^−^ = *P. stutzeri* no megaplasmid, pMP^+^ = *P. stutzeri* with pMPPla107. A = area and is measured in ten thousands of pixels. Images are representative of at least 3 replicates.

To determine if inhibitory agent is a common product produced by other bacteria, we also tested the supernatant of *E. coli* against DBL386 and DBL408. Zones of inhibition were not observed indicating that the supernatants of saturated *E. coli* do not contain inhibitory properties against DBL386 or DBL408 (Figure 7).

### Carriage of pMPPla107 Reduces Production of the Inhibitory Agent

We also hypothesized that if supernatants from various Pseudomonads inhibit the growth of pMPPla107 carrying strains, then megaplasmid acquisition could reduce production of the inhibitory agent. Our data indicate that the inhibitory agent activity is reduced in all tested supernatants collected from strains with pMPPla107 when compared to the supernatants of identical strains without pMPPla107 (Figure 7). Furthermore, this is not a product of differences in growth as all strains have reached saturation after 24 or 48hr of growth (Table 1). Although variation in the efficacy of the inhibitory phenotype occurs, all strains produce a similar diffuse inhibitory phenotype against strains carrying pMPPla107. Furthermore, acquisition of pMPPla107 results in reduced production of the compound across all species tested (Figure 7).

### The Inhibitory Agent is Bacteriostatic

Preliminary data in the lab suggested that the inhibitory agent is not bactericidal. To determine if the inhibitory agent has bactericidal or bacteriostatic activity we mixed filter sterilized supernatants containing the inhibitory agent with cells grown for four hours and stained the mixtures with SYTO9 and propidium iodide. SYTO9 is a green-fluorescent nucleic acid stain and will penetrate living cells while propidium iodide is a non-permeable nucleic acid stain that only stains cells with damaged membranes red-fluorescent. When mixing cells containing pMPPla107 with the inhibitory agent we found no observable cells with damaged membranes. These data suggest that the inhibitory agent is a growth inhibitor and does not kill cells at the naturally produced concentration.

## DISCUSSION

In this body of work we find that acquisition of pMPPla107 confers sensitivity to a non-proteinaceous small molecule compound that is resistant to both protease K and heat treatments(21, 27, 28). Furthermore, this inhibitory agent is conserved across Pseudomonads, is likely part of an essential or redundant pathway, and the concentration of the compound is positively correlated with the amount of time *Pseudomonas spp*. grow under laboratory conditions. These results suggest that the inhibitory agent is a commonly produced molecule among *Pseudomonas spp*. that may be a toxic secondary metabolite but that it is not produced by species like *E. coli*.

Due to the unknown nature of the *P. aeruginosa* supernatants on pMPPla107 containing strains, we optimized preferred media types and growth times to allow for clear and distinct overlays so that these methodologies can be used in future analyses of this phenotype. *P. fluorescens* and *P. putida* strains produced the inhibitory agent at higher levels when grown in KB media than in LB. Furthermore, KB was also the preferred media among all strains tested for obtaining clearer zones of inhibition when compared to LB. Therefore, nutrients found in KB medium may affect the metabolism of Pseudomonads to increase flux to pathways responsible for production of the inhibitory agent. Therefore, further analyses utilizing minimal media for plating overlays and the production of the inhibitory agent may give further insight into the metabolic pathways necessary for production and sensitivity characteristics of pMPPla107 and the inhibitory agent.

The inhibitory agent produced by *P. stutzeri* increases at a similar rate as growth in rich media, a characteristic shared by quorum sensing molecules. Quorum Sensing molecules are also known to recognize inter and intraspecies interactions and can induce several types of responses in bacteria including production of antibiotics(2, 29–33). All quorum sensing pathways are represented in the PA14 transposon mutant library and screening of this library revealed no single knockout in quorum sensing pathways that eliminated or reduced production of the inhibitory agent. Furthermore, the screened library did not reveal a single knockout, which eliminated production of the inhibitory agent. This can be explained by three possibilities: 1) the inhibitory agent is the product of redundant pathways requiring multiple knockouts, 2) one of the 335 genes without representation in the library produces the inhibitory agent (See Table 5 in Liberati et al.)(23), thus a knockout is not present in the library, or 3) the knockout mutation is present in one of the 95 mutants lost in testing. Therefore, the lack of inhibitory activity in *E. coli* supernatants against pMPPla107 strains could be due to an essential gene unique to *Pseudomonas spp*., which is not present in *E. coli*.

Although nutrient deprivation can elicit antagonistic stress responses in cells, this response is not typically induced in clonal populations (such as a pure culture) except in a case where population density and spread were shown to be regulated by active self-lysis via secreted proteins in low-nutrient environments(2, 8, 34). Here we report a set of different results that demonstrate an inhibitory agent isolated from a clonal population with rich nutritional resources inhibits the growth of nearly identical bacteria with the addition of pMPPla107, yet does not inhibit itself. One possibility for this outcome is that pMPPla107 carries a gene sensitive to the conserved inhibitory agent, which results in bacteriostatic growth inhibition. The inhibitory agent could also share a similar role to a previously mentioned self-lysing growth regulation system(34), where pMPPla107 is more susceptible to such a signal due to the burden and fitness costs it imposes on the cell. In this case the expectation would be that costs like decreased thermotolerances, decreased biofilm production, and increased antibiotic sensitivities previously found(21) would be correlated with sensitivity to the inhibitory agent. Cellular networks are complex and heavily regulated to maintain fitness, and the introduction of a megaplasmid carrying nearly a thousand genes could disrupt these pathways causing previously benign compounds to result in toxicity through over accumulation of a specific product or byproduct.

Addiction systems are small genetic elements that encode a stable toxin that can several cellular processes (e.g. transcription, translation, and ATP synthesis) and a less stable, but more abundant antitoxin(35). Plasmids commonly encode addiction systems as a means of self-preservation, as host cells will commonly attempt to cure plasmids without selective benefits. pMPPla107 carries multiple toxin/antitoxin systems that could is likely used for maintenance of the megaplasmid in microbial communities do to it’s low copy number(26, 36). Therefore, the inhibitory agent may interact with one of these cellular processes in conjunction with the toxins present on pMPPla107 resulting in additive negative effects on host cells causing growth inhibition. Whether this is a self-governing system being disrupted or truly antagonistic behaviors against pMPPla107 is still to be seen, but can be resolved by our ongoing attempts to identify the active inhibitory molecule through LC-MS and nuclear magnetic resonance analyses.

Acquisition of plasmids can contribute to several costs including protein interactions that disrupt cellular networks resulting in cytotoxic effects due to improper protein dosing, alteration to metabolic flux, or sequestration of RNA polymerases(12, 13, 37–39). Sensitization to the inhibitory agent by pMPPla107 offers yet another insight into cost associated HGT by demonstrating a direct cost to strains that have recently acquired pMPPla107 within a community. Given a pseudomonas community of cells bearing pMPPla107 and those lacking it, this inhibitory agent will be present within the community actively inhibiting growth of cells containing pMPPla107 and is therefore interesting that pMPPla107 conjugates efficiently within the lab and in nature(26, 36). However, we also demonstrate that acquisition of pMPPla107 reduces production of the inhibitory agent suggesting that perhaps this cost can be overcome with reduction of this compound coupled with high conjugation efficiency and retention.

Utilizing an interdisciplinary set of methods, we build upon our previous publication regarding a unique inhibitory phenotype(21) and find that pMPPla107 increases sensitivity to a small molecular growth inhibitory compound, is conserved across the pseudomonas phylogeny, and the gene responsible for its production is either redundant or essential. This suggests a highly conserved toxic compound that is yet another cost associated with pMPPla107 and a potential defense mechanism against this megaplasmid. Furthermore, we demonstrate through fluorescence microscopy that the molecule has bacteriostatic properties, indicating that at the naturally produced concentrations the inhibitory agent is a growth inhibitor to pseudomonas cells with pMPPla107. The data presented here provides foundational knowledge of a system to pursue in depth analyses centered on how horizontally transferred elements can alter microbial interactions, result in antibacterial costs, and potentially be utilized as a mobile sensitivity system.

**Figure 8:**
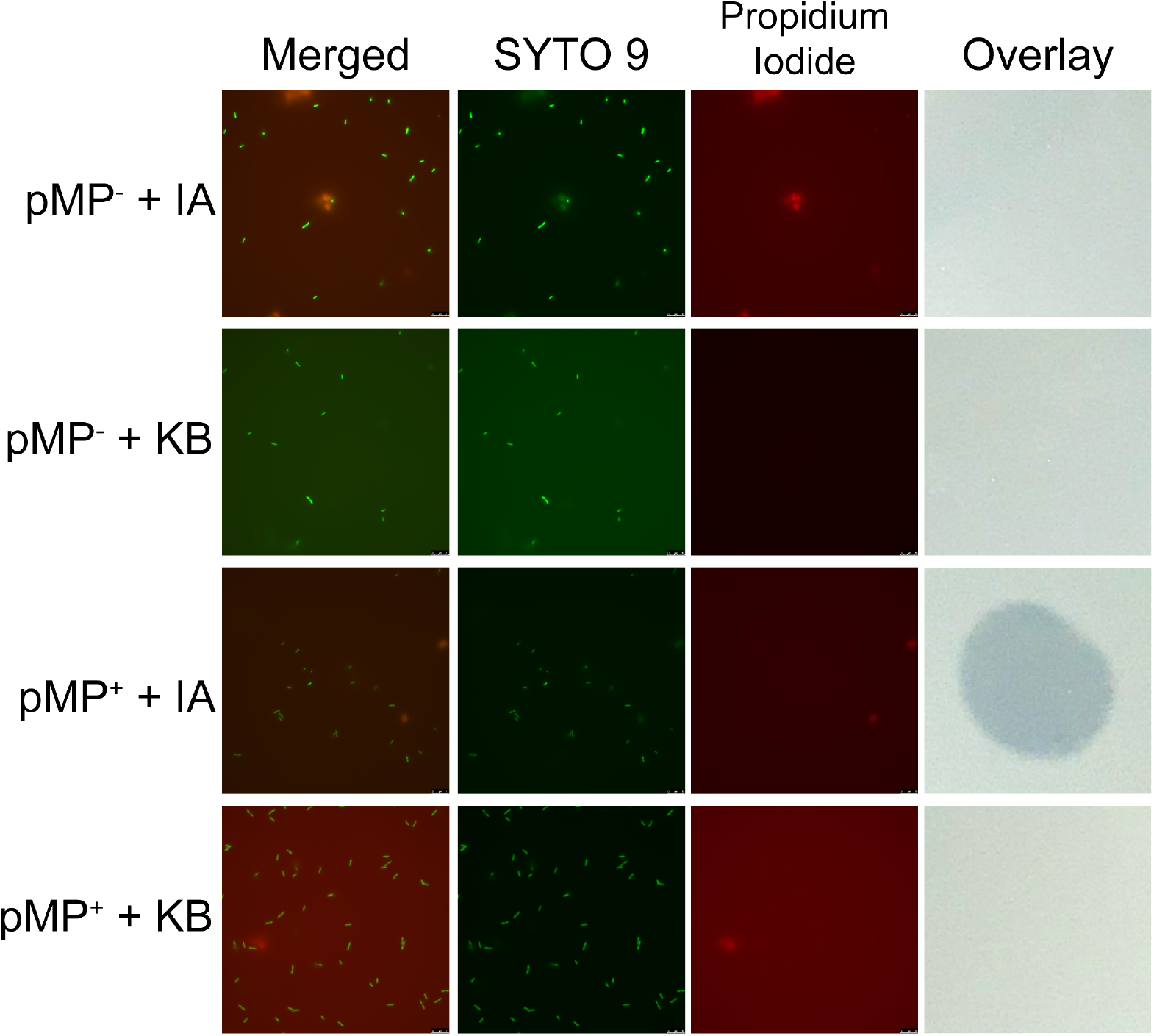
Fluorescent staining suggests the inhibitory agent has a bacteriostatic effect on cells. LIVE/DEAD staining of cells mixed 1:1 with either the inhibitory agent or KB (negative control) did not reveal dead cells (red) stained by propidium iodide; instead all cells were living (green) stained by SYTO9. pMP^−^ = *P. stutzeri* without a megaplasmid (DBL386), pMP^+^ = *P. stutzeri* with pMPPla107 (DBL408), IA = Inhibitory Agent, KB = King’s medium B. Images are representative of at least 3 replicates. Images from a corresponding overlay are present demonstrating the expected inhibitory agent phenotype in these samples.

## Supplemental Figures

**Supplemental Figure 1:**
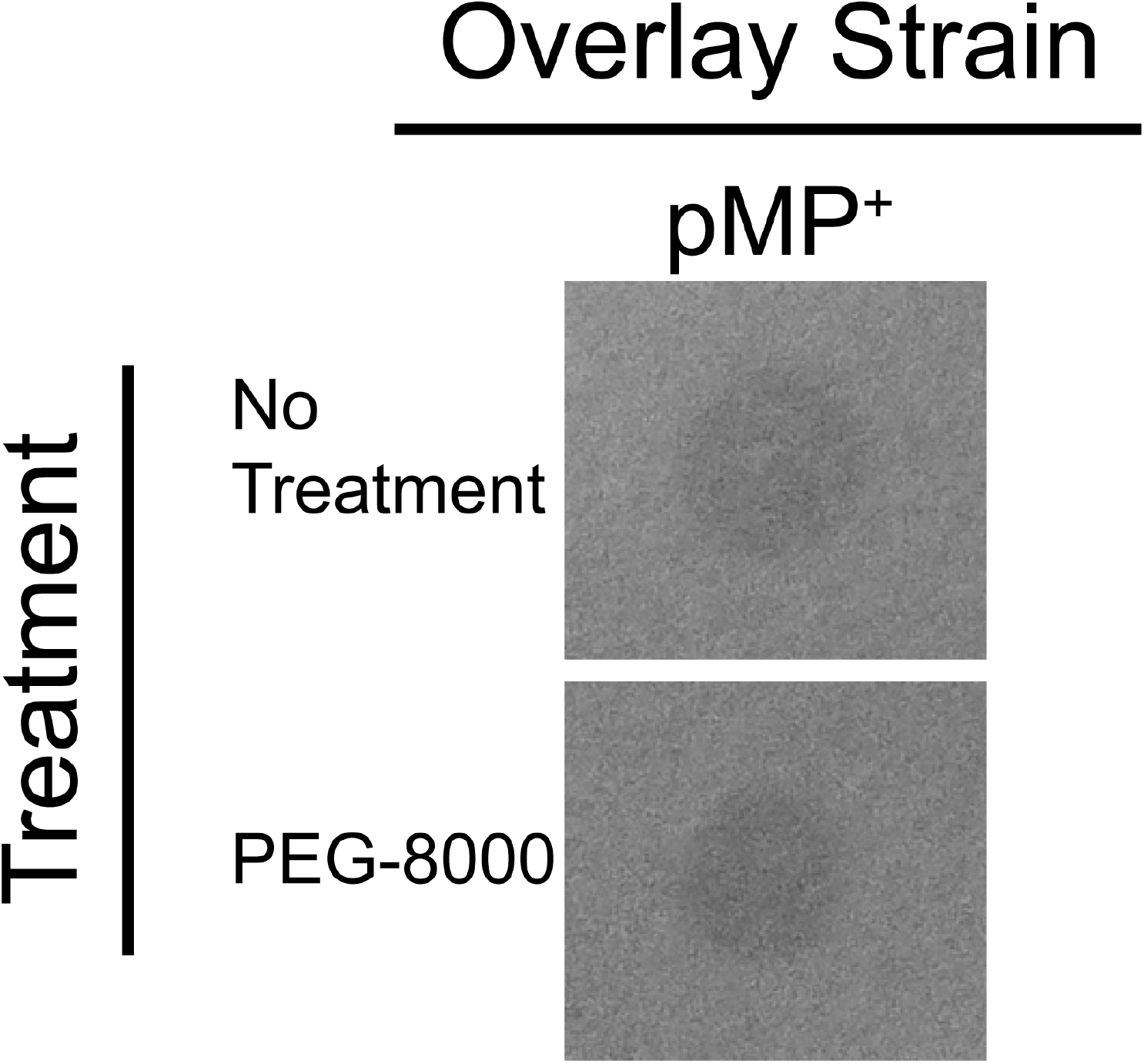
Protein precipitation with PEG-8000 does not concentrate the inhibitory agent activity. The supernatants were collected from *P. aeruginosa* grown for 24hrs in LB. Supernatants were filter sterilized and pulled down with PEG-8000 or not pulled down. 10μL spots were of these samples were placed on an overlay of *P. stutzeri* with pMPPla107. Overlay strains were grown for four hours in KB before plating. All images are representative of 3 biological replicates.

**Supplemental Figure 2:**
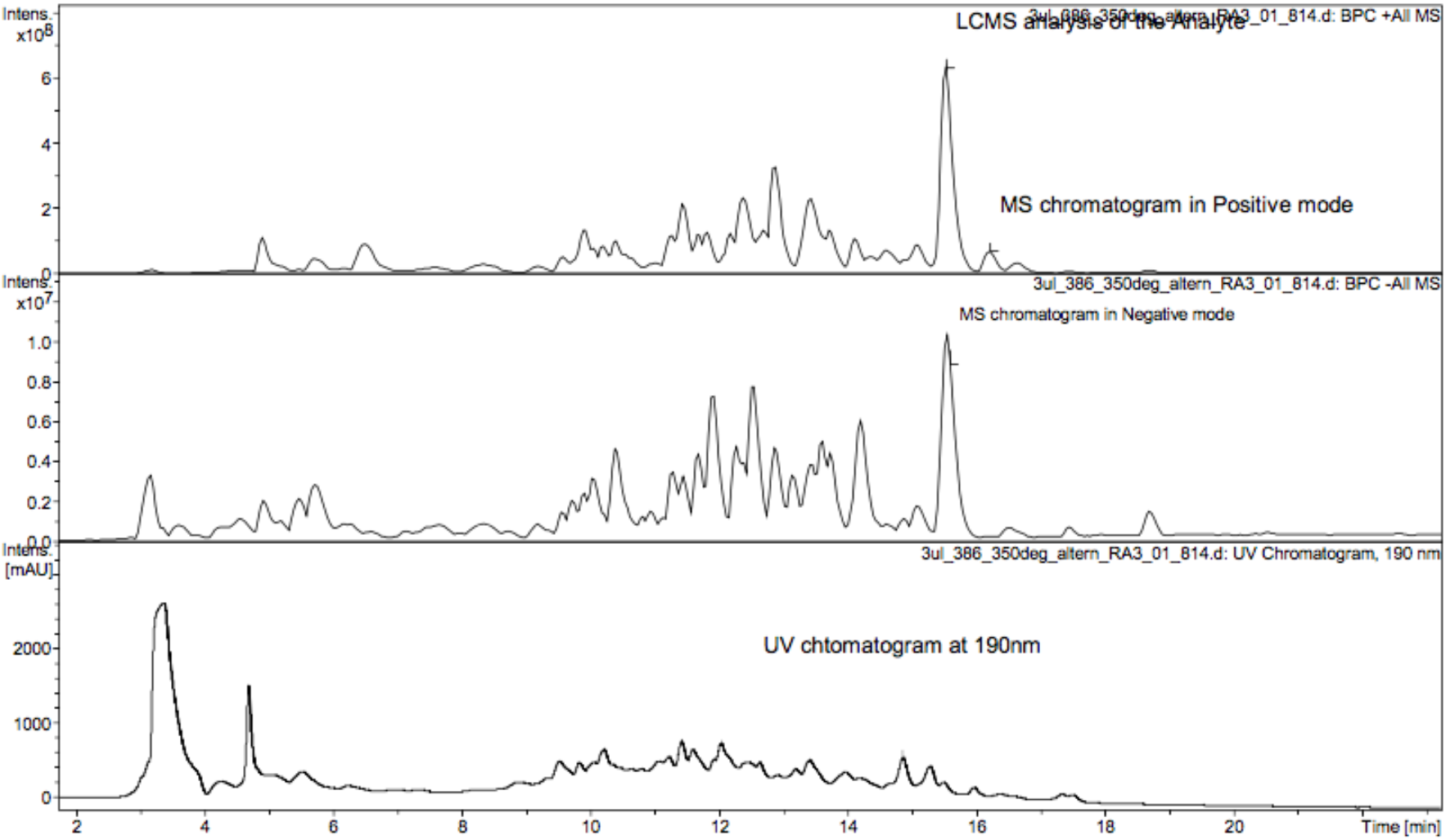
LCMS analysis of the inhibitory agent. LCMS analysis. LC-MS analysis of the analyte was performed via amaZon with the sample LC separation via the ThermoFisher Scientific UltiMate 3000 HPLC system indicates that the supernatant sample is highly complex due to many peaks in MS and LC chromatograms.

**Supplemental Figure 3:**
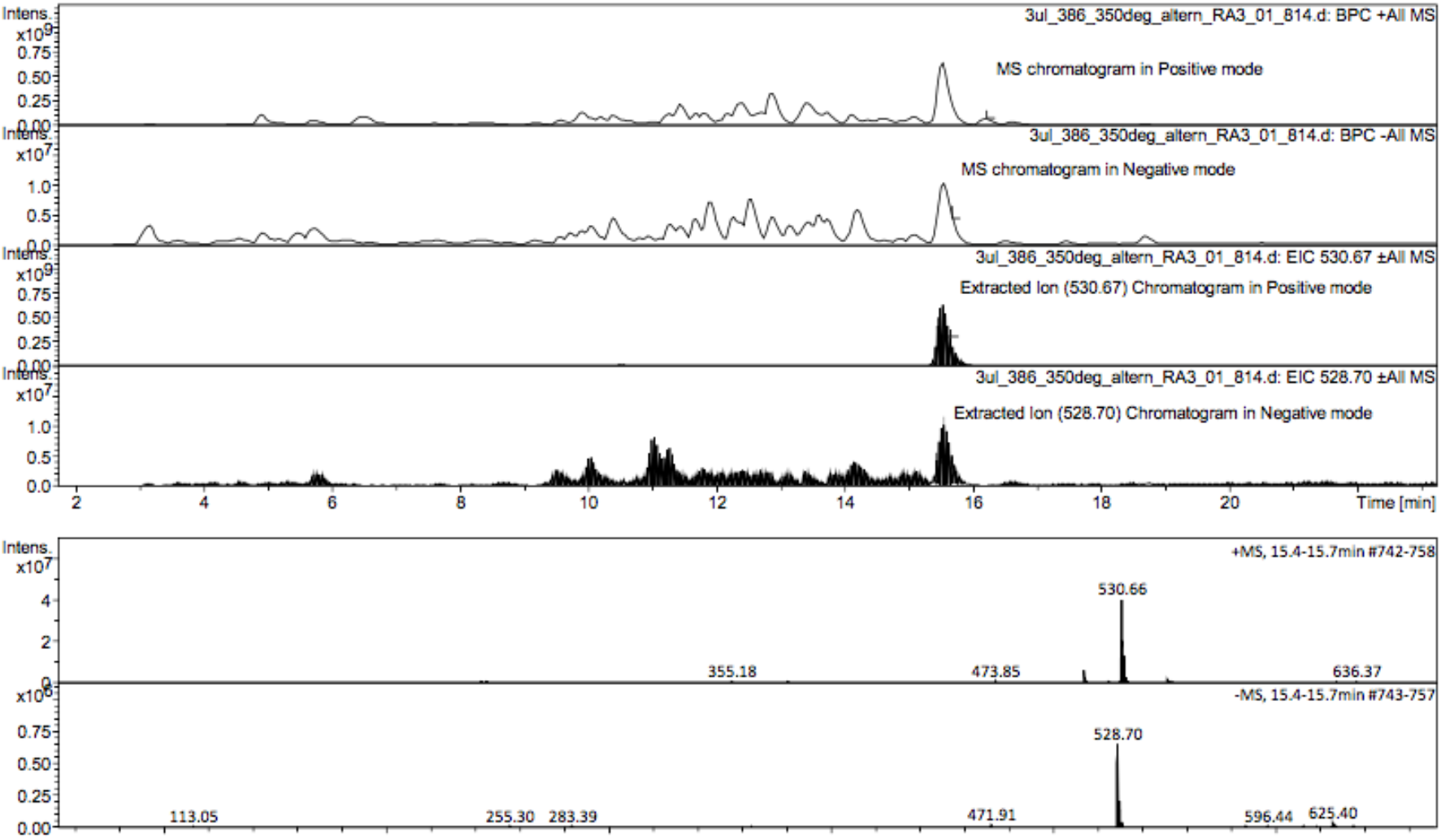
Extracted Ion Chromatograms for 530.72 and 528.70. Extraction of these masses defines the retention time for the inhibitory agent at 15.5min allowing for multiple fraction collections and submission to NMR analysis.

**Supplemental Figure 4:**
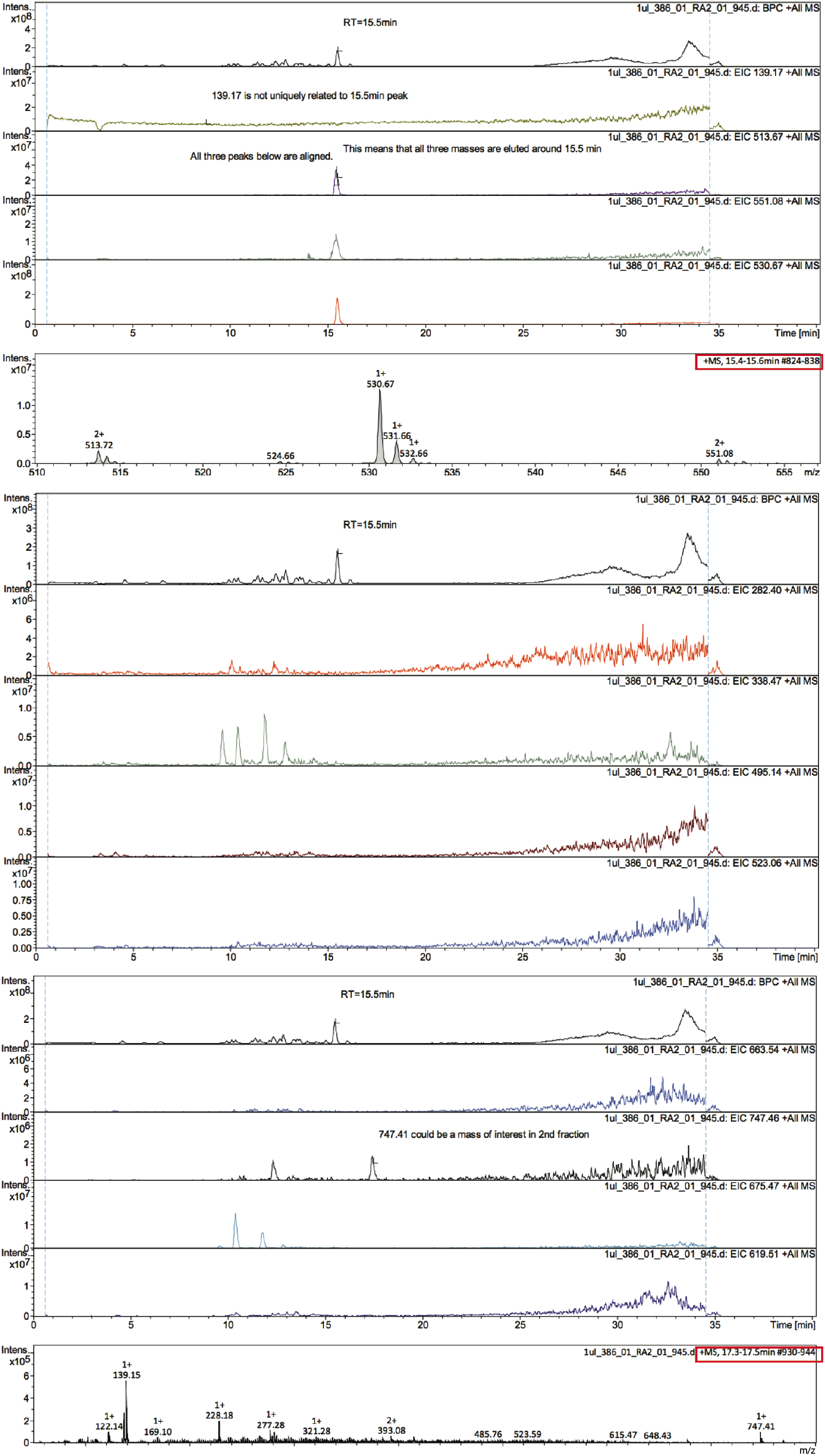
UV Chromatograms and Mass Spectrums of Outstanding Peaks. UV chromatograms of potential masses of interest reveal two time points with distinct peaks of interest in fractions 1 and 2 (15.4-15.6min and 17.3-17.5min). Specific masses for each chromatogram are indicated as EIC XXX.XX in the top right corner. Their corresponding mass spectrums are also presented and their time points are boxed in red. Masses 513.72 and 551.08 are doubly charged (2+) and demonstrate low intensity peaks while 530.67 is singly charged with a more intense peak.

